# Deriving shape-based features for *C. elegans* locomotion using dimensionality reduction methods

**DOI:** 10.1101/054379

**Authors:** Bertalan Gyenes, André E.X. Brown

**Affiliations:** MRC Clinical Sciences Centre, Du Cane Road, London, UK; Institute of Clinical Sciences, Faculty of Medicine, Imperial College London, Du Cane Road, London, UK; Department of Mathematics, Imperial College London, UK

## Abstract

High-throughput analysis of animal behavior has become a reality with the advance of recording technology, leading to large high-dimensional data sets. This dimensionality can sometimes be reduced while still retaining relevant information. In the case of the nematode worm *Caenorhabditis elegans*, more than 90% of the shape variance can be captured using just four principal components. However, it remains unclear if other methods can achieve a more compact representation or contribute further biological insight to worm locomotion. Here we take a data-driven approach to worm shape analysis using independent component analysis (ICA), non-negative matrix factorization (NMF), a cosine series, and jPCA and confirm that the dimensionality of worm shape space is close to four. Projecting worm shapes onto the bases derived using each method gives interpretable features ranging from head movements to tail oscillation. We use these as a comparison method to find differences between the wild type N2 worms and various mutants. The different bases provide complementary views of worm behavior and we expect that closer examination of the time series of projected amplitudes will lead to new results in the future.

## Introduction

Animal behavior is a high-dimensional problem since each joint in vertebrates and each independent muscle in invertebrates introduces new degrees of freedom. This makes it challenging to provide comprehensive and quantitative descriptions of behavior even in small animals like the nematode worm *Caenorhabditis elegans* (Gomez-Marin et al., 2014). Traditional ethology methods have focused on observer-defined categories to reduce behavioral dimensionality but automated imaging and data analysis tools have made it possible to extract more complete records of an animal’s behavior (Anderson and Perona, 2014; Chen and Engert, 2014; Gouvêa et al., 2014; Machado et al., 2015; Ohyama et al., 2013; Ramdya et al., 2015). From these data, lower-dimensional representations can then be identified using unsupervised learning algorithms. Dimensionality reduction can be achieved using a variety of different methods. Each emphasizes different aspects of the underlying behavior and it is not clear which of these will be the most informative in advance or in fact what behavioral feature each corresponds to in contrast to observer-defined categories. However, the assumptions and limitations of each automated approach are made explicit in the algorithm and they can be compared quantitatively on a common data set.

The nematode worm *C. elegans* is a useful model to test different dimensionality reduction methods. *C. elegans* moves by propagating bending waves along its body and, when confined to the surface of an agar plate, this motion occurs in two dimensions, making it possible to capture its behavior using a single camera. Previous work on *C. elegans* body shape using principal component analysis (PCA) has shown that the effective dimensionality of worm locomotion is low, as there are correlations between bends along different parts of the body (Stephens et al., 2008). Trajectories through the lower dimensional space defined by the principal components can be used to classify different genotypes and explain certain behaviors both in *C. elegans* and in the larvae of *Drosophila melanogaster* (Brown et al., 2013; Stephens et al., 2011; Szigeti et al., 2015).

Here we revisit the question of how to represent worm shape space by using four different dimensionality reduction methods. As each of these methods has different objectives, the resulting dimensions highlight different aspects of *C. elegans* shape space. We analyze these differences using features derived from the methods and compare the behavior of mutant worms.

## Methods

### Data

The dataset used in the analysis was collected and described previously (Yemini et al., 2013). It contains 9964 videos of single worms moving freely on an agar plate for 15 minutes (after a 30- minute-long acclimatization period). 335 different genotypes were analyzed including the N2 lab strain. We used the angle representation of the worm (Fig. 1B-C) with a mean of zero except for non-negative matrix factorization (NMF) where all values were made positive by adding a constant (a requirement of the method). 10 N2 trajectories were picked randomly from a collection of 100 as the training set for jPCA. To obtain the variance of the basis shapes, we resampled the same collection 100 times obtaining 10 trajectories each time.

**Figure 1.**
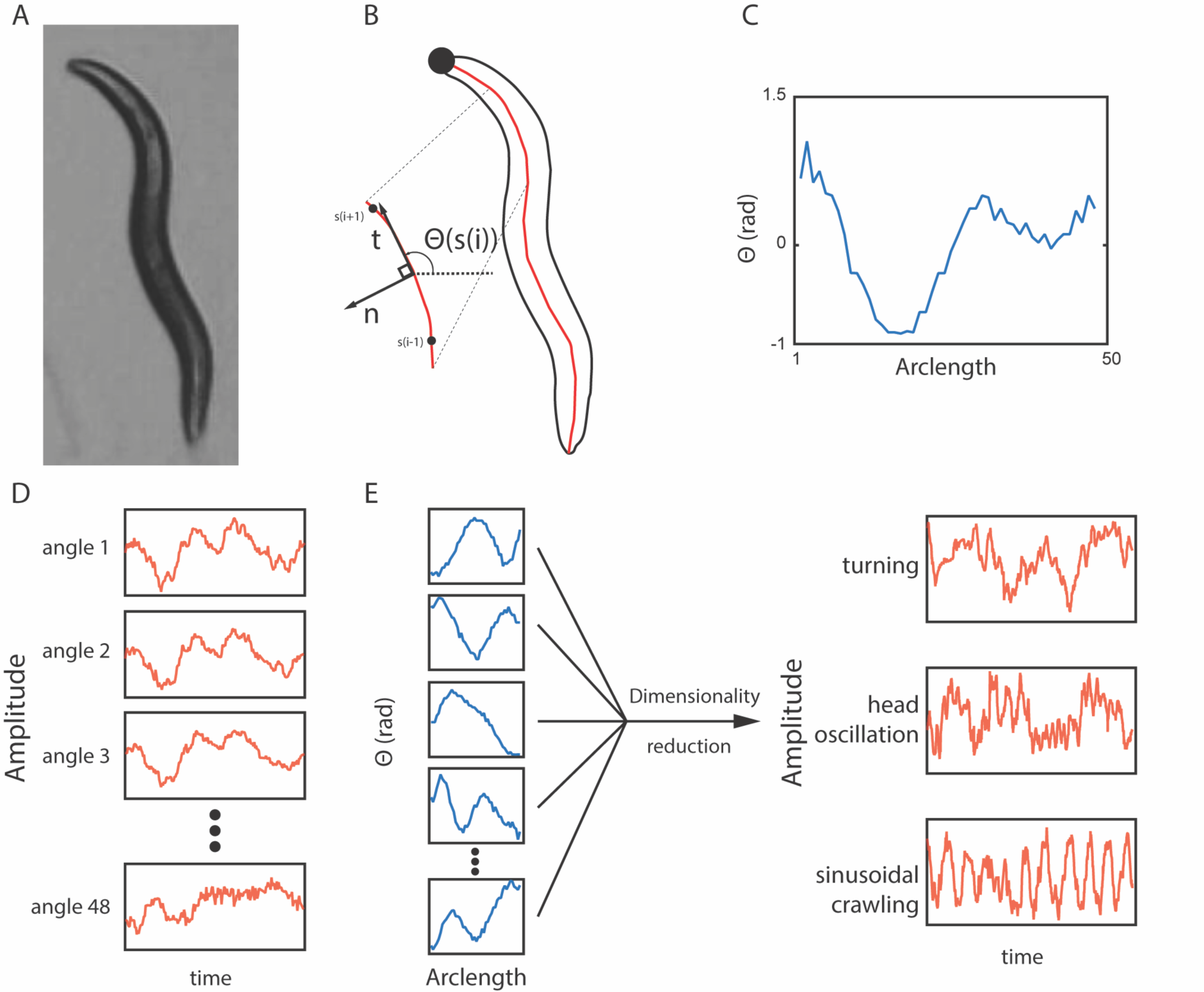
(A) A typical frame of a worm under the tracking microscope. (B)-(C) The outline and the curve through the center of the worm. The angle between neighboring points along the centerline is plotted from the tip of the head (s = 1) to the end of the tail (s = 48). (D) As the worm moves, the value of each angle changes, but each subsequent angle provides little additional information because they are highly correlated with each other. (E) Dimensionality reduction methods can reveal more biologically meaningful time-series variables.

### Dimensionality reduction

A training set of 3000 N2 shapes was picked randomly from a collection of 12600 for ICA and NMF. To obtain the variance of the basis shapes, we resampled the same collection 100 times obtaining 3000 N2 shapes each time. For analysis, a testing set of 3000 N2 shapes was projected onto each basis shape to retrieve the corresponding amplitudes. To ensure that all of the mutants were represented in the test between PCA and the sinusoidal basis shapes, we sampled 1 shape from each of the 9964 recordings in the dataset. Each worm shape was reconstructed using either four principal components or the sinusoidal basis and the squared difference between the reconstructed and original shapes were determined in each case. PCA and NMF were conducted using built-in functions of MATLAB, while freely available methods were used for ICA (http://research.ics.aalto.fi/ica/fastica/, (Hyvarinen, 1999)) and jPCA (Churchland et al., 2012). We used the deflation approach and the power law nonlinearity as the parameters for ICA, but we find that our results are robust to different parameters, as well. The sinusoidal basis shapes were defined to be cosine waves,

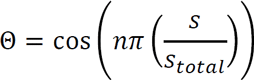

where *s* is the arclength, *s_total_* is the total arclength, and *n* is an integer from 1 to the number of basis shapes used.

### Mutant comparisons

We projected the entire dataset onto the NMF and jPCA basis shapes (derived from the N2 wild type training set) to obtain the projected amplitudes for each worm at each time. We then took the mean absolute value of each projected amplitude as a simple feature characterizing each worm’s average shape. For jPCA, we measured the mean amplitude of the anterior oscillation in each individual, i.e. 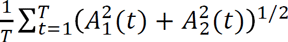, where *A_1_* and *A_2_* are the projections onto the first two eigenshapes, and *t* is the frame number. These were compared between each genotype and the wild type (N2) using a Mann-Whitney U-test. Bonferroni correction was used to control for multiple comparisons.

### Worm maintenance and recordings

As previously described (Yemini et al., 2011), worms were maintained under standard conditions on NGM plates with OP50 as food source at 22 °C. Induced reversal experiments were carried out as described in (Alkema et al., 2005).

## Results

### Independent component analysis refines features derived from PCA

Independent component analysis (ICA) minimizes the statistical dependence of the components in multivariate signals as compared with PCA that minimizes the projection error. This means that ICA can remove noise and separate artifacts from the data (Hyvärinen and Oja, 2000), while PCA focuses on reducing the unexplained variance with successive components.

We find that ICA returns four basis shapes that are reminiscent of the ones obtained using PCA (Fig. 2A-B), but the projected amplitudes of full worm trajectories show clear differences. This is consistent across resamplings and different parameters. The two PCA eigenshapes shown in Figure 2C have previously been described as forming an approximate quadrature pair (Stephens et al., 2008). Therefore, the travelling wave that worms form during crawling locomotion is encoded as phase-shifted oscillations in these modes. Histograms of projections onto the first two basis shapes averaged over multiple worms are shown in Fig. 2. A ring structure suggesting oscillatory behavior is clearly present during forward locomotion (Fig. 2C, top row) with both methods, although the probability distribution is less constant along the ring using PCA compared to ICA. When all the data are plotted including turns and dwelling, the probability distribution becomes more uniform, especially for PCA. This suggests that ICA returns modes that isolate the crawling wave more completely from other aspects of the shape dynamics compared to PCA.

**Figure 2.**
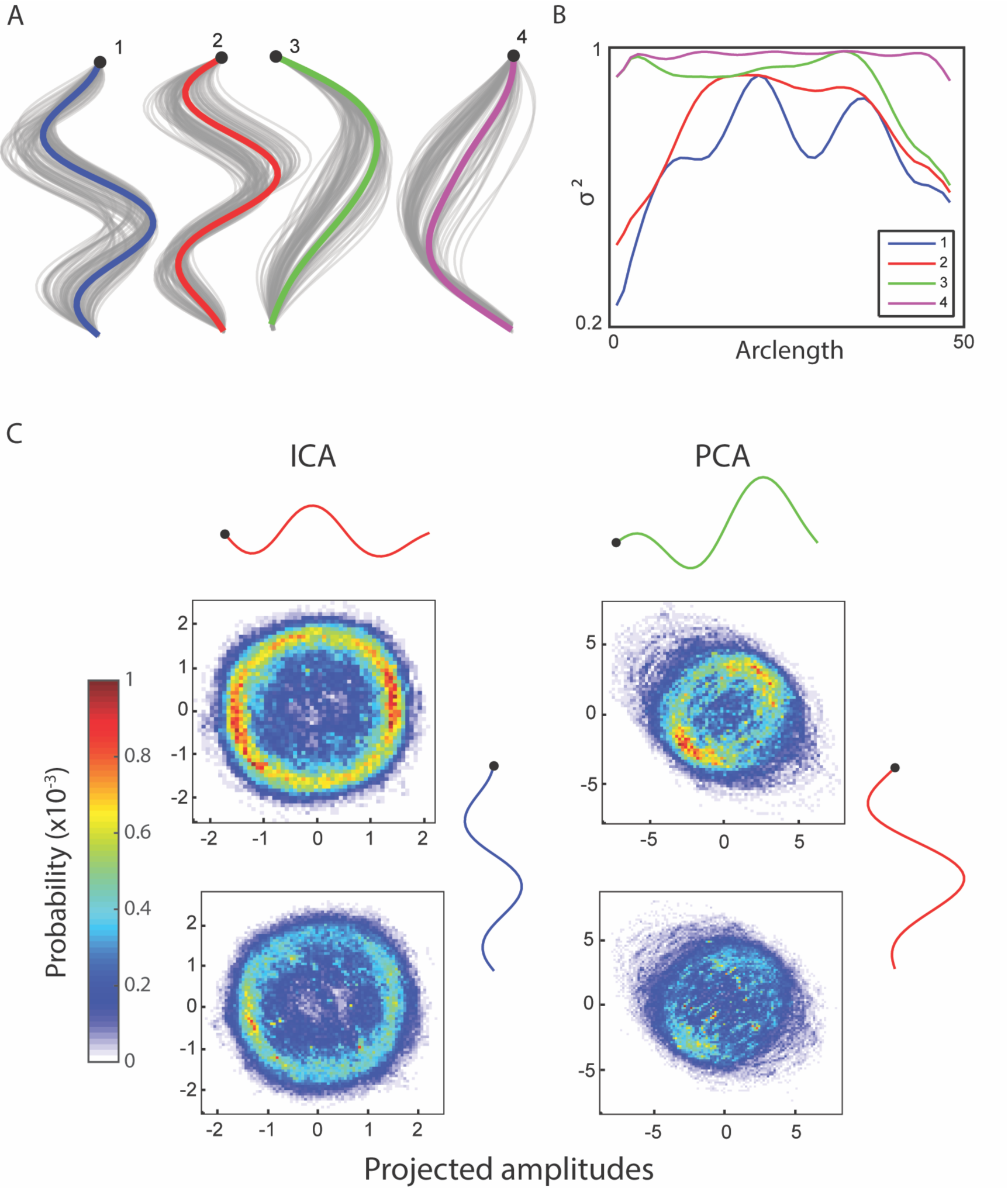
(A) Independent component analysis returns four basis shapes that explain 97.6% of the variance in the dataset. The graph shows an x-y coordinate representation of the modes with the resampled basis shapes in grey. (B) The fraction of the variance explained along the worm by including an increasing number of basis shapes suggests that the modes can each explain a different part of the worm well. (C) Bivariate histograms for the amplitudes of basis shapes (wild type worm, 15 minutes, frame rate: 30 Hz). Top row: forward locomotion only, bottom row: all data. Basis shapes 1 and 2 from ICA form a ring in both cases (especially clear when only the forward locomotion is counted), suggesting an oscillatory behavior between them. Similarly, two basis shapes from principal component analysis are known to explain an oscillatory behavior, but they also include other information, as evidenced by a lack of clear, continuous ring in their histograms.

### Worm body segments are individually defined by non-negative matrix factorization

Non-negative matrix factorization is a commonly used method in computer vision and data clustering (Lee and Seung, 1999). In contrast to other methods that are more focused on returning a combination of the original variables as the reduced dimensions, NMF finds a parts-based representation. In the shape dataset, this means that each of the basis shapes is going to be good at explaining a particular segment of the worm and the corresponding amplitude will directly correlate with the size of displacement in that segment. Running the algorithm returns a set of basis shapes that indeed divides the worm into 5 approximately equally spaced segments (Fig. 3A-B) corresponding to the head, neck, midbody, hip and tail regions.

**Figure 3.**
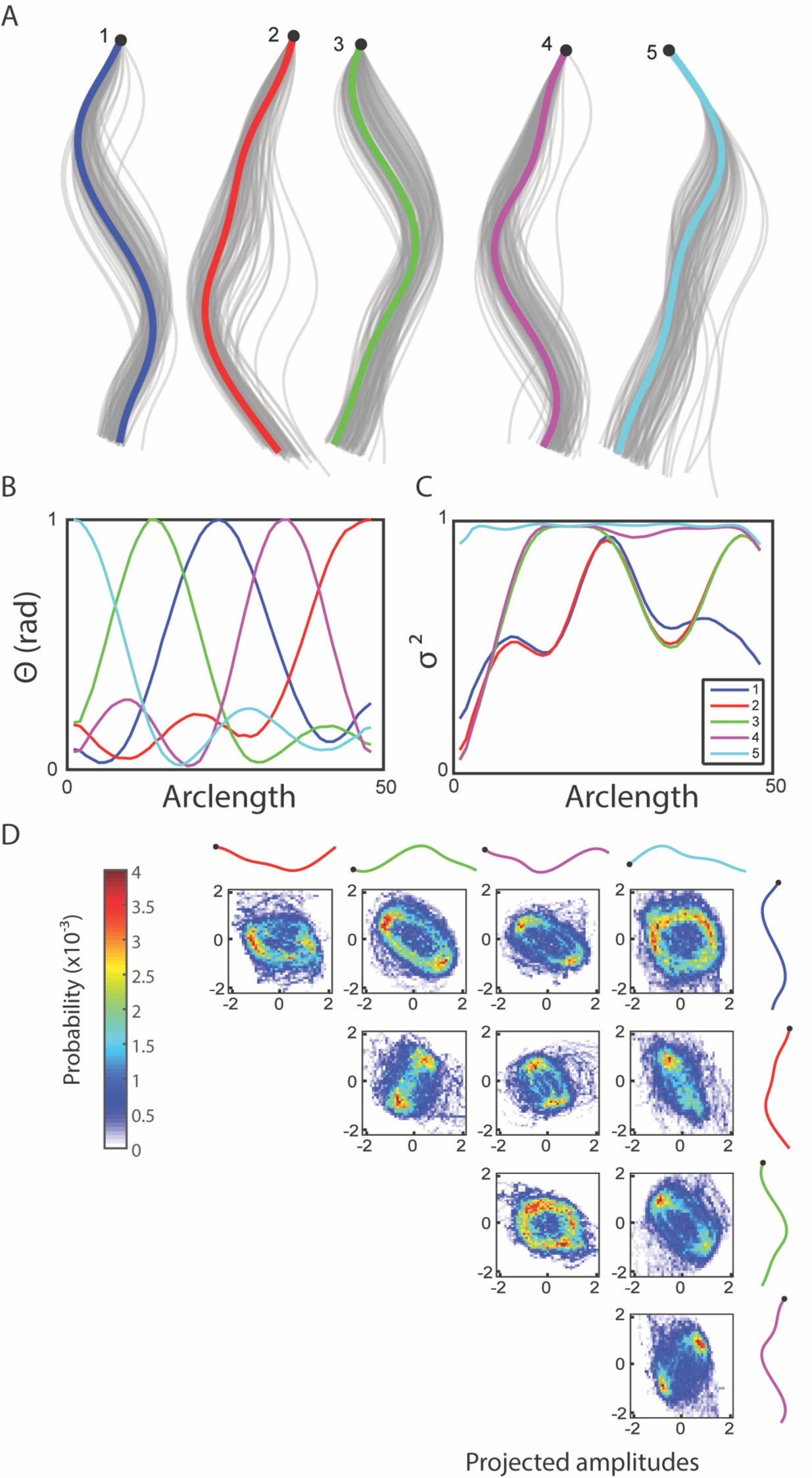
(A) Non-negative matrix factorization returns five basis shapes that explain 97.6% of the variance in the angle data. The graph shows an x-y coordinate representation of the modes with the resampled basis shapes in grey. (B) Angle representation of the basis shapes in (A) (legend in (C)). (C) The fraction of the variance explained along the worm by including an increasing number of basis shapes suggests that the modes can each explain a different part of the worm well, in this case localized to the five major segments of the worm. (D) Bivariate histograms for the amplitudes of basis shapes (wild type worm, 15 minutes, frame rate: 30 Hz). Basis shapes 3 and 4, and 1 and 5 both form incomplete rings, suggesting a more diffuse representation of the oscillatory sinusoidal crawling behavior using NMF.

We compared the NMF segment features (mean absolute projected amplitudes) across all 335 genotypes in the database using the basis shapes derived from the training set of wild type N2 shapes. This set of basis shapes captures 97.6% of the variance in N2 and 97.1% in mutants. At least one feature was significantly different compared to the wild type N2 strain in 172 genotypes (significance level: 0.01, Bonferroni corrected Mann-Whitney U-test) (Fig. 4). The results confirm earlier research: for instance the mutant *snf-6* is known to have exaggerated head movements (Kim et al. 2004). Most behavioral studies have not focused on describing the locomotion phenotype in detail, as this is often difficult to do by eye. However, NMF can provide testable hypotheses on the location of effect in a novel way, for instance with regards to the mode of action of the gene *nlp-1.* The lack of this neuropeptide is known to increase the turning rate of the worm via modulating the AIA neurons, but it is not obvious how these two are linked, as these neurons are highly interconnected with other neurons (Chalasani et al., 2010). We find that *nlp-1* mutants show an increase in the amplitude projected onto the mode that corresponds to the head, while there is no significant difference along other parts of the body (Fig. 4). Such a localized response could help constrain hypotheses for AIA function by focusing on neural circuits that modulate head muscles. NMF can also help in discerning phenotypes that may be masked by more obvious effects. An example of this is the *egg-5* mutant that has severe developmental problems during the oocyte phase while still in the parent worm (Parry et al. 2009). The increased movement in the hip and the tail of the worm (Fig. 4) could be due to a decrease of eggshell on the eggs inside the gonads, making them more flexible and less restrictive for the worm.

**Figure 4.**
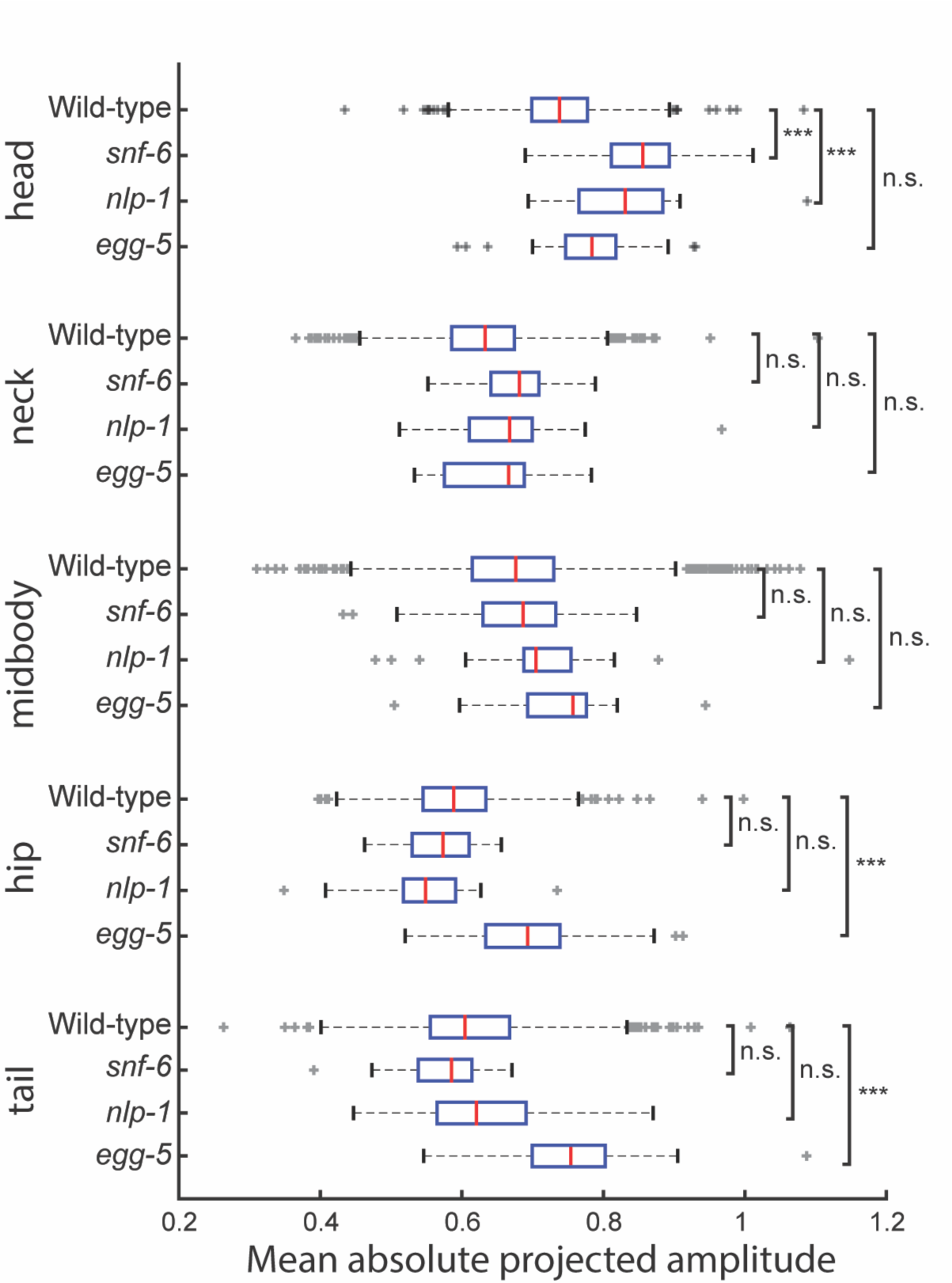
The mean of the absolute projected amplitudes corresponding to each basis shape from non negative matrix factorization is taken for individual worms of four different genotypes. (wild type N2: n = 1303, *snf-6:* n = 43, *nlp-1:* n = 22, *egg-5:* n = 23) *snf-6* and *nlp-1* worms have significantly increased head motion, but normal movement in the rest of their body in terms of magnitude (p_adj_(snf 6) = 3.13 × 10^−14^, p_adj_(nlp-1) = 6.83 × 10^−4^), while the opposite can be observed in *egg-5* mutants (padj(hip) = 8.19 × 10^−5^, padj(tail) = 2.48 × 10^−6^).

### Fourier cosine series captures 97% of variance across mutants

Data-driven dimensionality reduction methods are inherently dependent on the dataset used to train them, meaning that the basis shapes produced will be different if a different training set is used. If the training set is large enough, variation will be small, but if only a small number of trajectories are available for a given condition then the derived shapes could vary significantly from sample to sample. Using a set of pre-determined basis shapes would avoid this issue, but to be useful they must explain most of the shape space variance across different individuals. Given the sinuous set of basis shapes derived using both PCA and ICA, we defined a Fourier cosine series as a set of basis shapes and tested if it could capture worm shapes compactly (Fig. 5A-B). The first four basis shapes of the cosine series captured 96.9% of the variance across the mutant shape test set (Fig. 5C). While the cosine series performs significantly worse than PCA (p=2.49*10^−11^, t-test), the difference is small (the top four PCA components capture 97.1% of the variance) and may be negligible for some applications. Using a set of analytically defined modes may prove useful in theoretical applications.

**Figure 5.**
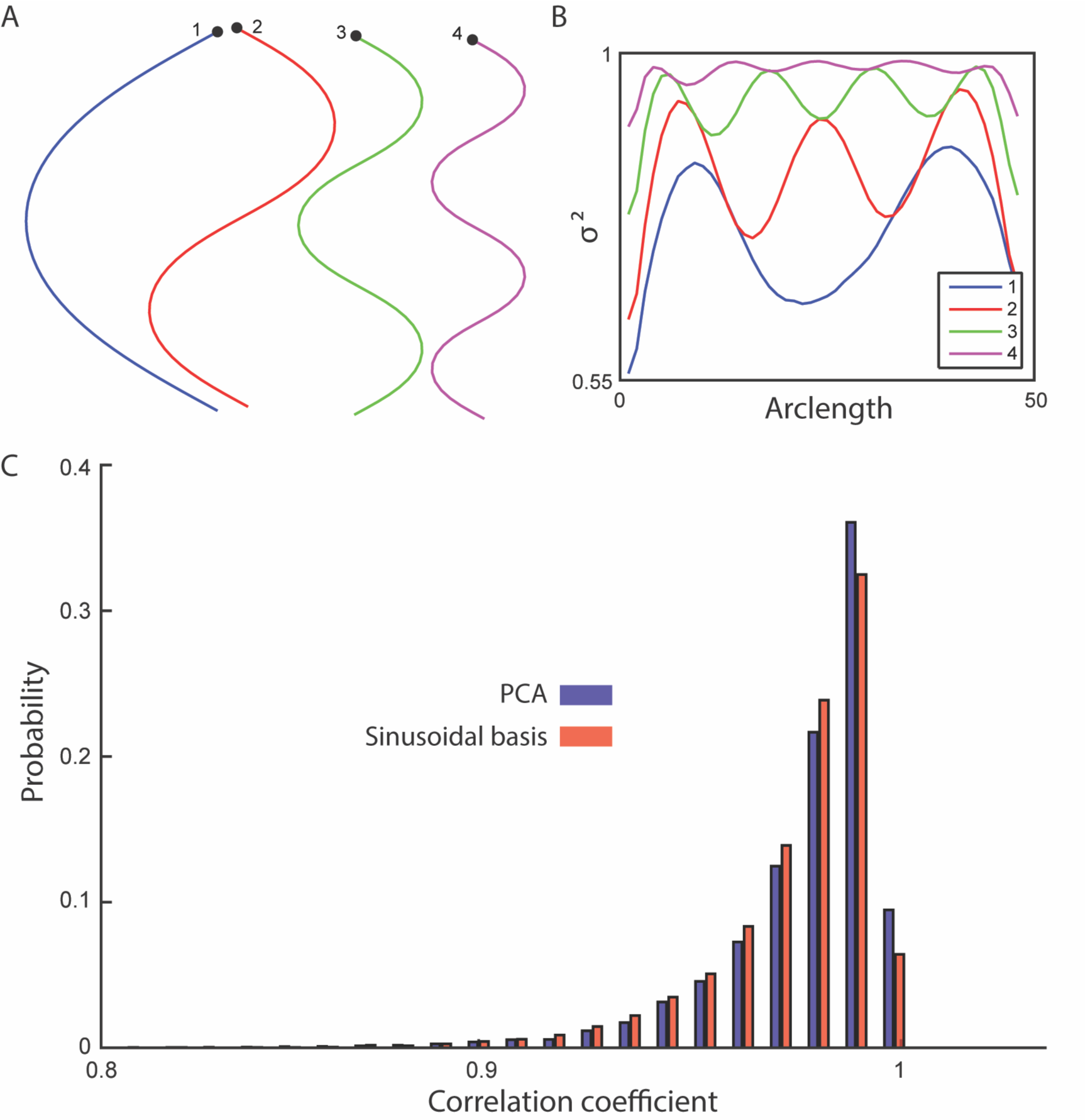
(A) A cosine series was used to generate four basis shapes with increasing frequency. The corresponding x-y representations are shown. (B) The fraction of the variance explained along the worm by including an increasing number of basis shapes. (C) The shapes in the testing set were reconstructed using the four sinusoidal basis shapes and the top four modes of principal component analysis. The histogram of the correlation coefficients (between the reconstructed and the original shapes) suggests a significant, but small difference between the sinusoidal analysis (96.9%) and the data-driven approach (97.1%) (t-test, p = 2.49 × 10^−11^).

### Body oscillations are described by jPCA

The methods considered above are time independent: they only take into account the distribution of shapes. In contrast, jPCA uses time series trajectories of worm motion, maps the shape space with PCA and then reorients these components to identify components that show strong oscillations (Churchland et al., 2012). Using this method on wild type (N2) trajectories leads to three pairs of components, each pair corresponding to a segment along the body of the worm (Fig. 6A-B). The components are ordered according to the strength of the oscillation detected, indicating that the oscillation produced during locomotion decreases in strength from head to tail on average.

**Figure 6.**
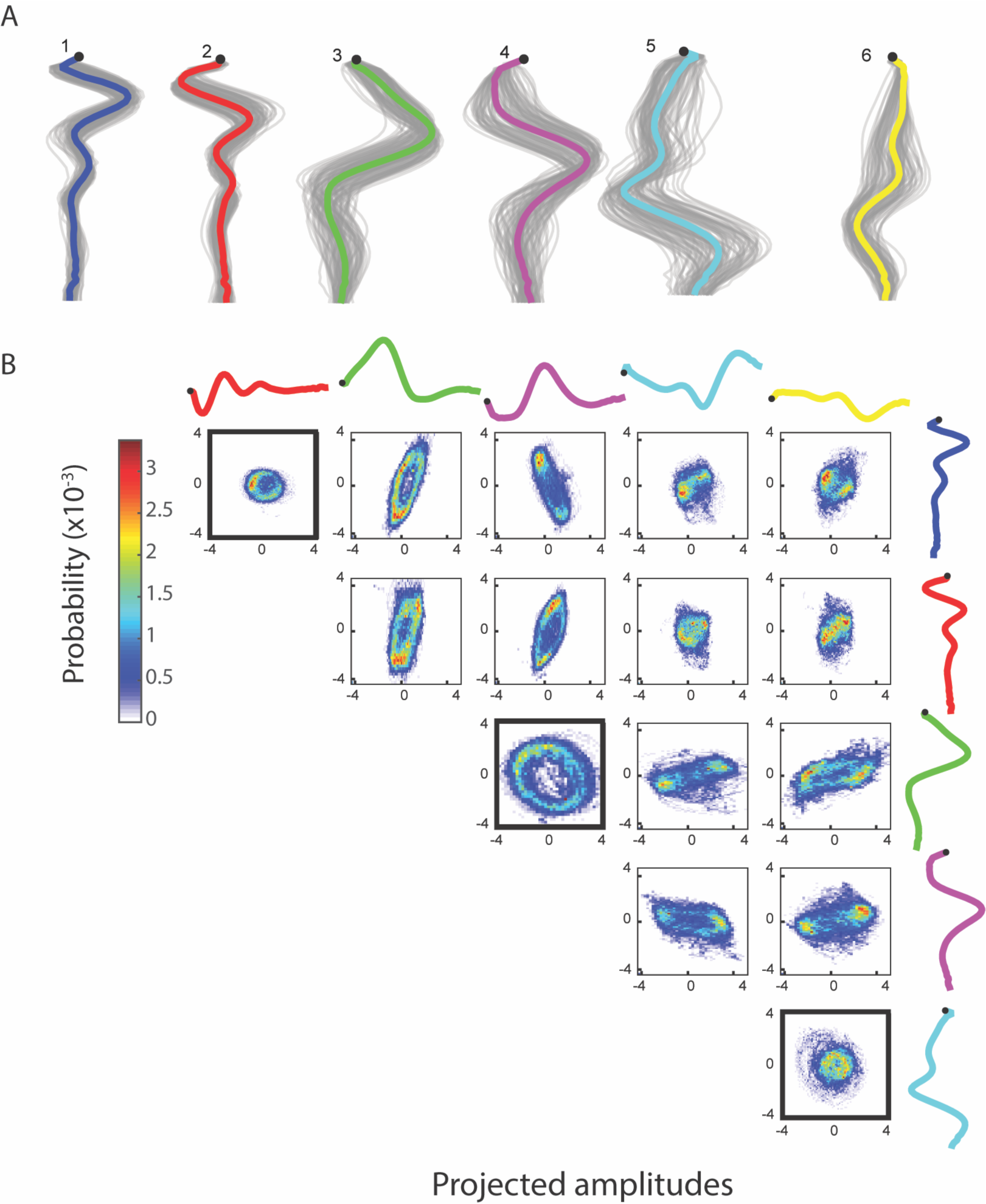
(A) jPCA is run with 12 components (top 6 shown here). The graph shows an x-y coordinate representation of the modes with the resampled basis shapes in grey. (B) Bivariate histograms for the amplitudes of basis shapes (wild type worm, 15 minutes, frame rate: 30 Hz). Basis shapes 1 and 2, 3 and 4, and 5 and 6 all form rings, suggesting an oscillatory behavior between them and independent sinusoidal waves in the corresponding parts of the body.

Worms have different movement patterns during reversals as opposed to forward motion. We analyzed different mutants to see if there is any difference compared to wild type N2 by looking at the anterior body oscillation, a behavior that was the most rotationally robust in the dataset. Similarly to NMF, there is a large number of mutants (168, significance level: 0.01) significantly different compared to wild type N2 in the size of the anterior oscillation. Two examples are shown in Fig. 7. We found that the wild type worm reduces the size of its anterior body oscillation during spontaneous reversals, prompting us to consider whether this feature was sensitive to the head tip oscillation of the worm, as this is known to be suppressed during reversal (Alkema et al., 2005). However, the anterior oscillation detected by jPCA is not suppressed during touch-evoked reversals (Fig. 7). We also looked at *tdc-1(n3419)* mutants, which have been reported to maintain their head tip oscillation during touch-evoked reversals (Alkema et al., 2005). As with N2, we do not detect a change in jPCA anterior oscillation in *tdc-1(n3419)* touch-evoked reversals, but we do find that the magnitude of the oscillation is lower in *tdc-1* during spontaneous forward locomotion. This suggests that the jPCA anterior oscillation is not the same as the small oscillation that worms exhibit at the very tip of their heads. Despite this, the jPCA anterior oscillation does show a difference between spontaneous and touch-evoked reversals: both wild type and *tdc-1* worms show a stronger anterior body oscillation during touch-evoked reversals (Fig. 7). Finally, we also found that *egg-5* mutants fail to suppress their anterior body oscillation during reversal, even though they behave normally during forward locomotion.

**Figure 7.**
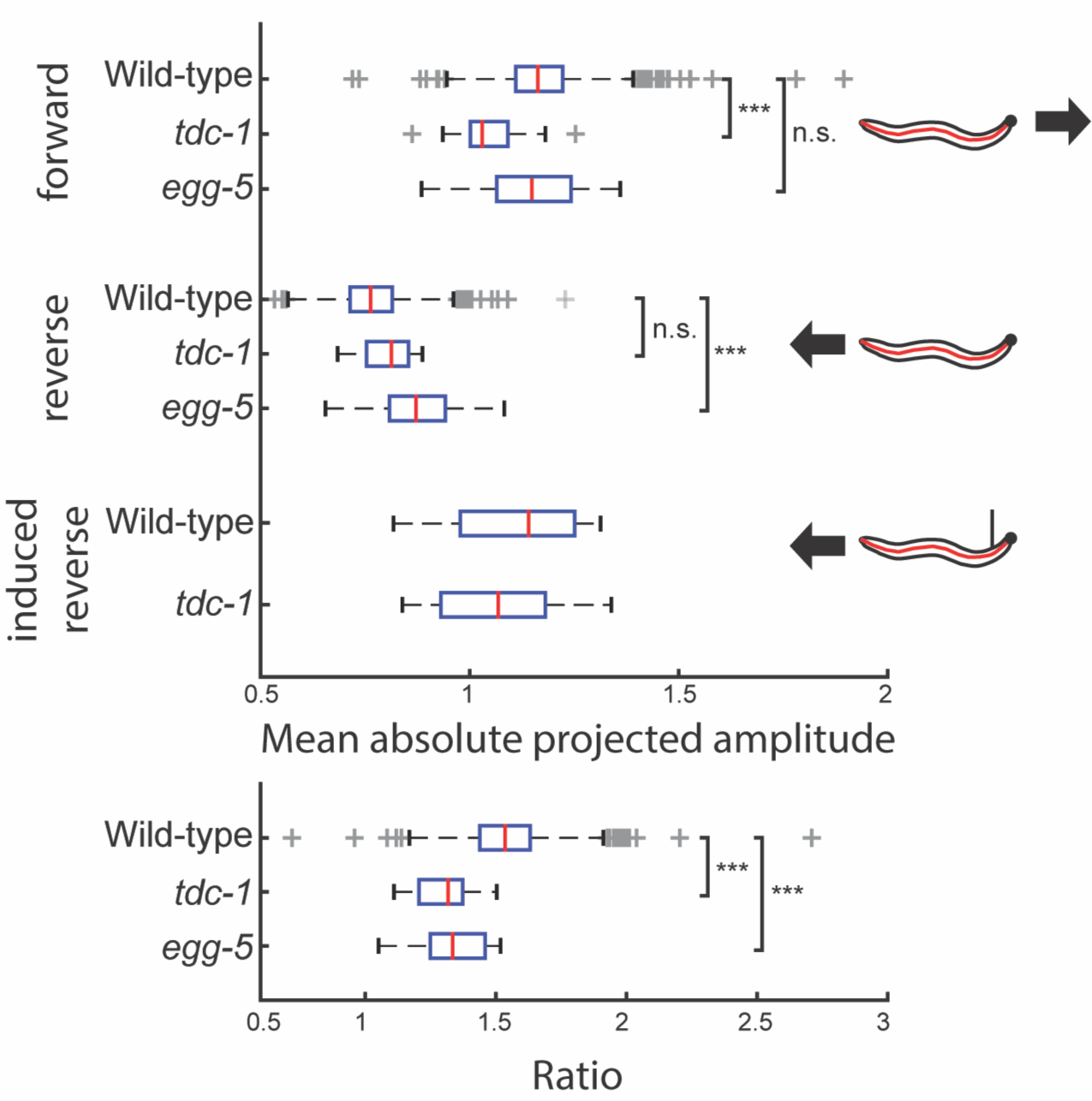
The amplitude of the jPCA anterior oscillation is measured for individual worms of three different genotypes during forward locomotion and reversals. (wild type N2: n = 1303, *tdc-1:* n = 19, *egg-5:* n = 23) *tdc-1* has significantly reduced head oscillation during forward locomotion, but suppresses it during reversals to the same magnitude as wild types (p_adj_(tdc-1) = 4.80 × 10^−5^), while the opposite can be observed in *egg-5* mutants (p_adj_(egg-5) = 3.71 × 10^−4^). During touch-evoked reversals, head oscillation is reduced in both wild type N2 and *tdc-1* worms. Both have a significantly smaller ratio (forward/spontaneous reversal) than wild type (p_adj_(tdc-1) = 3.73 × 10^−6^, p_adj_(egg-5) = 6.69 × 10^−6^).

## Discussion

We used four different dimensionality reduction methods to obtain a number of new features that can be used to describe different groups of worms. The new features are straightforward to use and show interpretable differences between mutants.

We found that none of the methods returned a more compact representation of the *C. elegans* shape space compared to PCA, confirming the previous lower-bound dimensionality of four for the worm shape space. However, different projections provide different kinds of information, for instance the intuitive joint-like representation of postural dynamics through non-negative matrix factorization or the full-body oscillations from independent component analysis. In addition, ICA clearly defines two sets of basis shapes (1 and 2; 3 and 4) corresponding to two waves with different frequencies, suggesting a possible representation of worm behavior as a superposition of two fundamental oscillations. The set of sinusoidal basis shapes provides an analytically defined set of shapes that could be used across experiments and labs to make results more directly comparable since they generalize well across mutant strains. jPCA contributes an interesting insight into the dynamic oscillation patterns of the worm body. This pattern could be consistent with a central pattern generator in the head producing an oscillation that becomes less coherent as it propagates down the worm (Gjorgjieva et al., 2014).

Behavior is a dynamic process often involving shifts between different states, single events and cyclic episodes. Our new features also change over time, but this was not taken into account when we interrogated the database. Instead, we used the magnitude averaged over the entire recording that reflects the general shape of the worm, which was sufficient to detect many significant differences. However, thorough time series analysis would likely reveal more details about the locomotion trajectories. Oscillations are ubiquitous in all four bases, but each feature also has a rich dynamical profile with different properties and comparison between these has the potential to provide different and complementary information. One example could be the characterization of the spontaneous switch between the feeding states of the worm. *C. elegans* has been reported to have three different behavioral states (roaming, dwelling and quiescence) that are influenced by food availability and nutritional status (You et al., 2008). The states are traditionally defined by instantaneous midbody speed when using automatic tracking, but this is known to have its limits when trying to find well-defined states (Ben Arous et al., 2009; Fujiwara et al., 2002; Gallagher et al., 2013). Shape has been useful for detecting lethargus, a different quiescent state that has a specific posture associated with it (Iwanir et al., 2013; Nelson and Raizen, 2013). The new shape features could provide further insight into shape differences that characterize different behavioral states. At the same time, some bases may be better suited than others for defining predictors of single events such as omega turns, and description of periodic behaviors like reversals.

Worm behavior has often been described using states defined by the experimenter. Using recording equipment and automated feature extraction was initially conceived to help with the process of group assignment and definition (Baek et al., 2002; de Bono and Bargmann, 1998), and this has been extended with unsupervised methods to detect patterns in worm locomotion (Brown et al., 2013; Schwarz et al., 2015). As we have shown here, the basis used for representing shape can reveal different aspects of behavior and provide new avenues for the future development of behavior classification and analysis methods.

